## Supplementary figures and images for "Deficiency of IQCH causes male infertility in humans and mice"

### Figure 1-source data-blots.tif

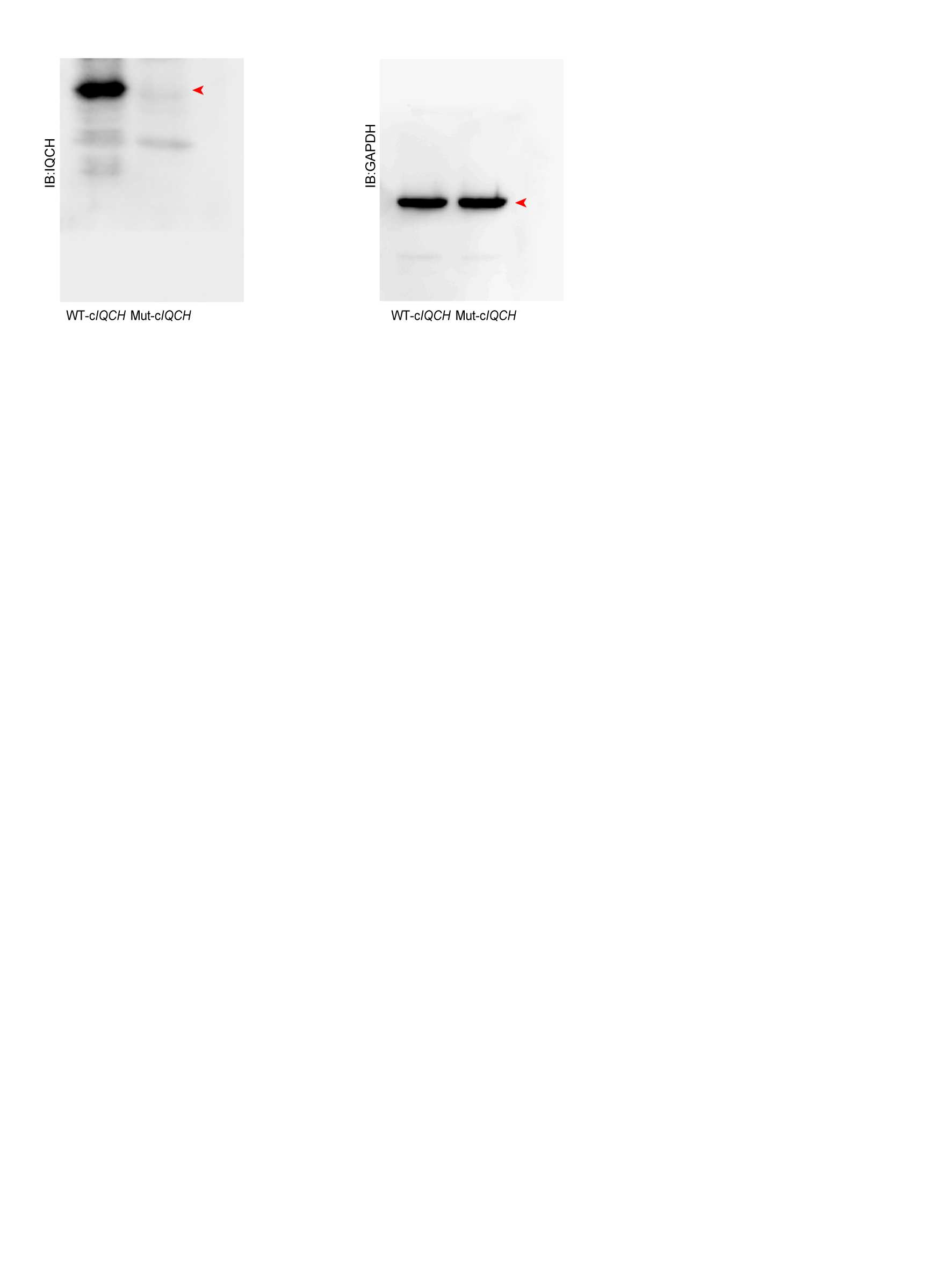

### Figure 1-source data-gels.tif

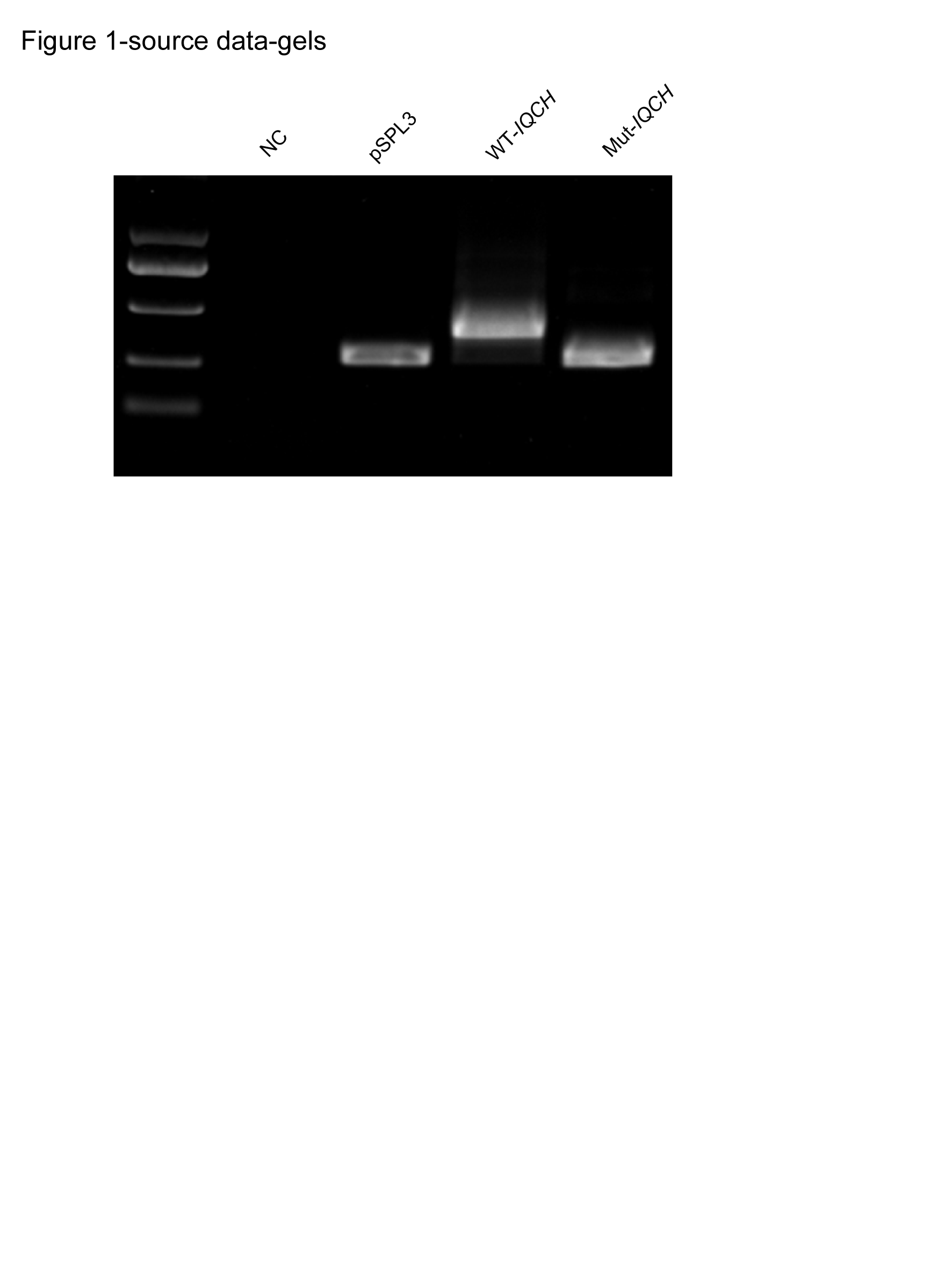

### Figure 3í¬figure supplement 1-source data-gels.tif

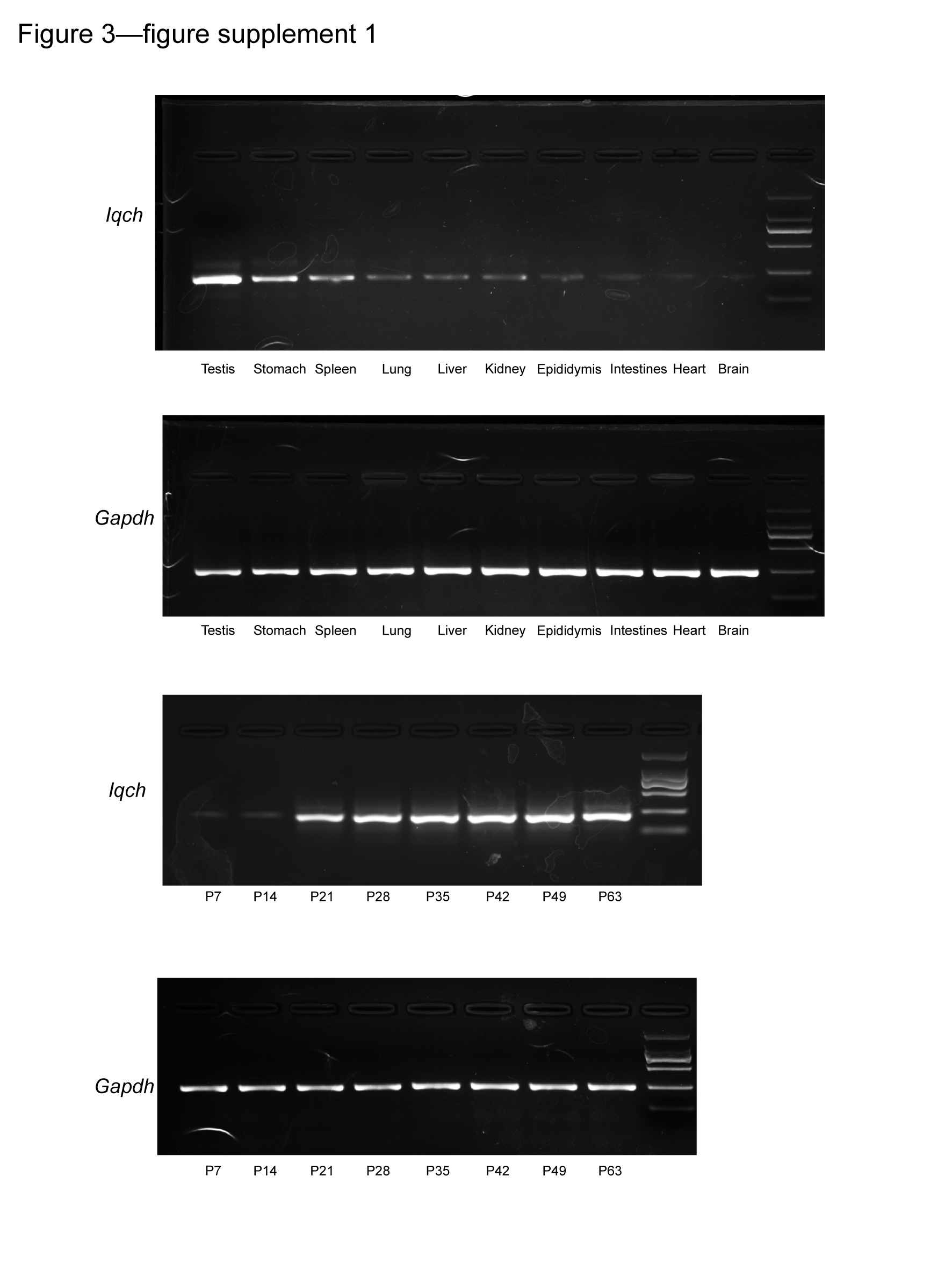

### Figure 3í¬figure supplement 2-source data-blots.tif

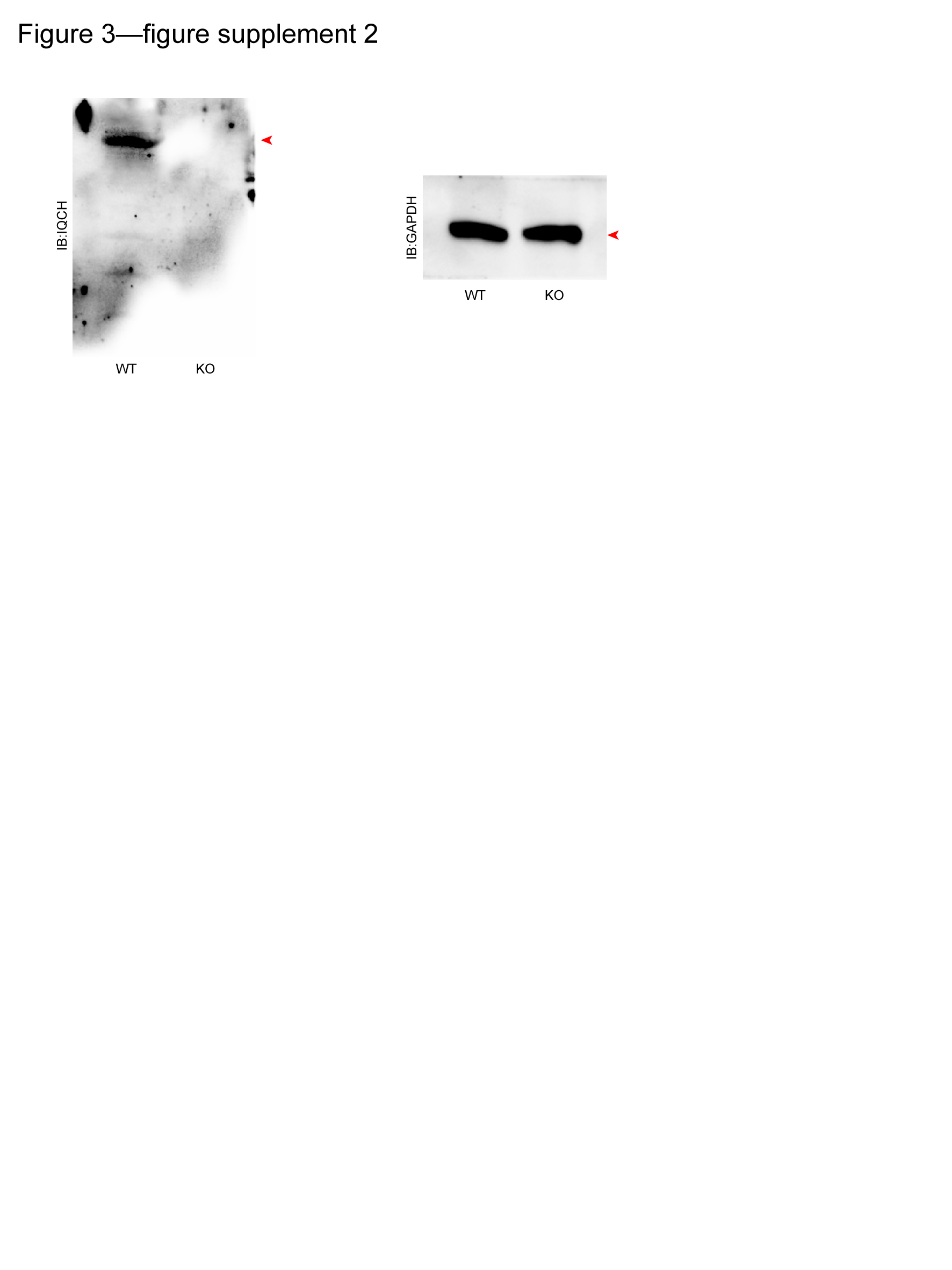

### Figure 3í¬figure supplement 2-source data-gels.tif

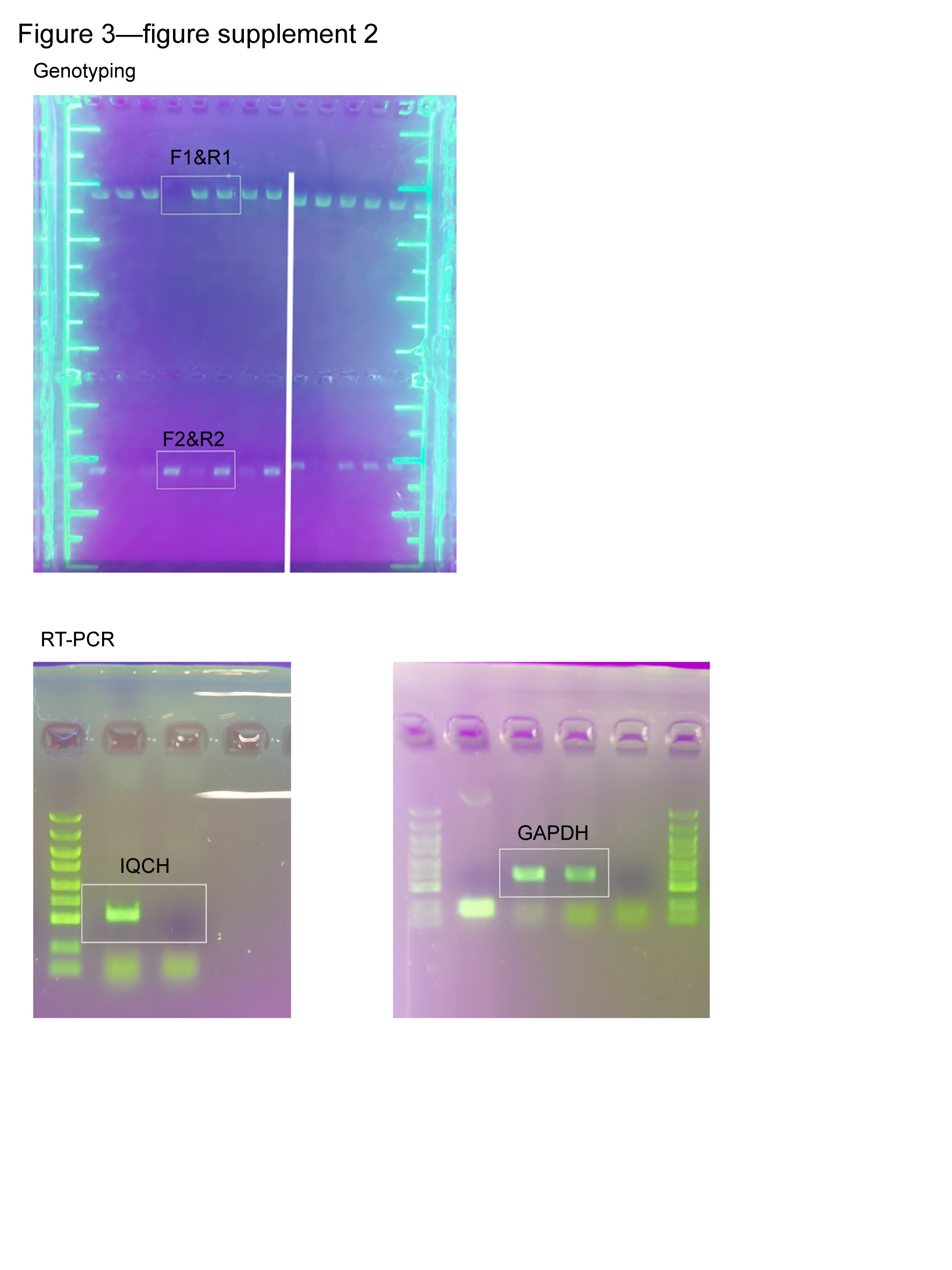

### Figure 6A-source data.tif

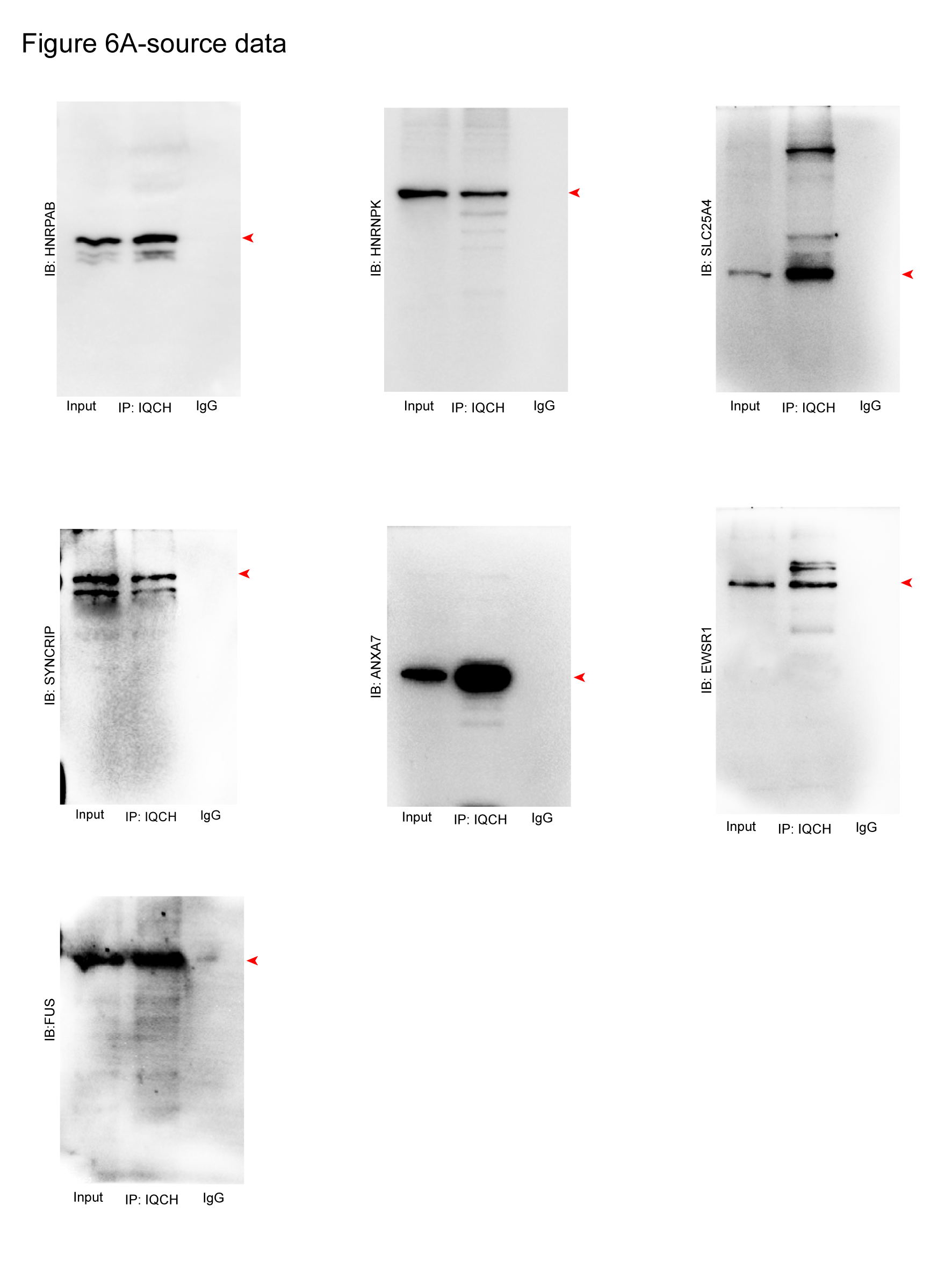

### Figure 6B-source data.tif

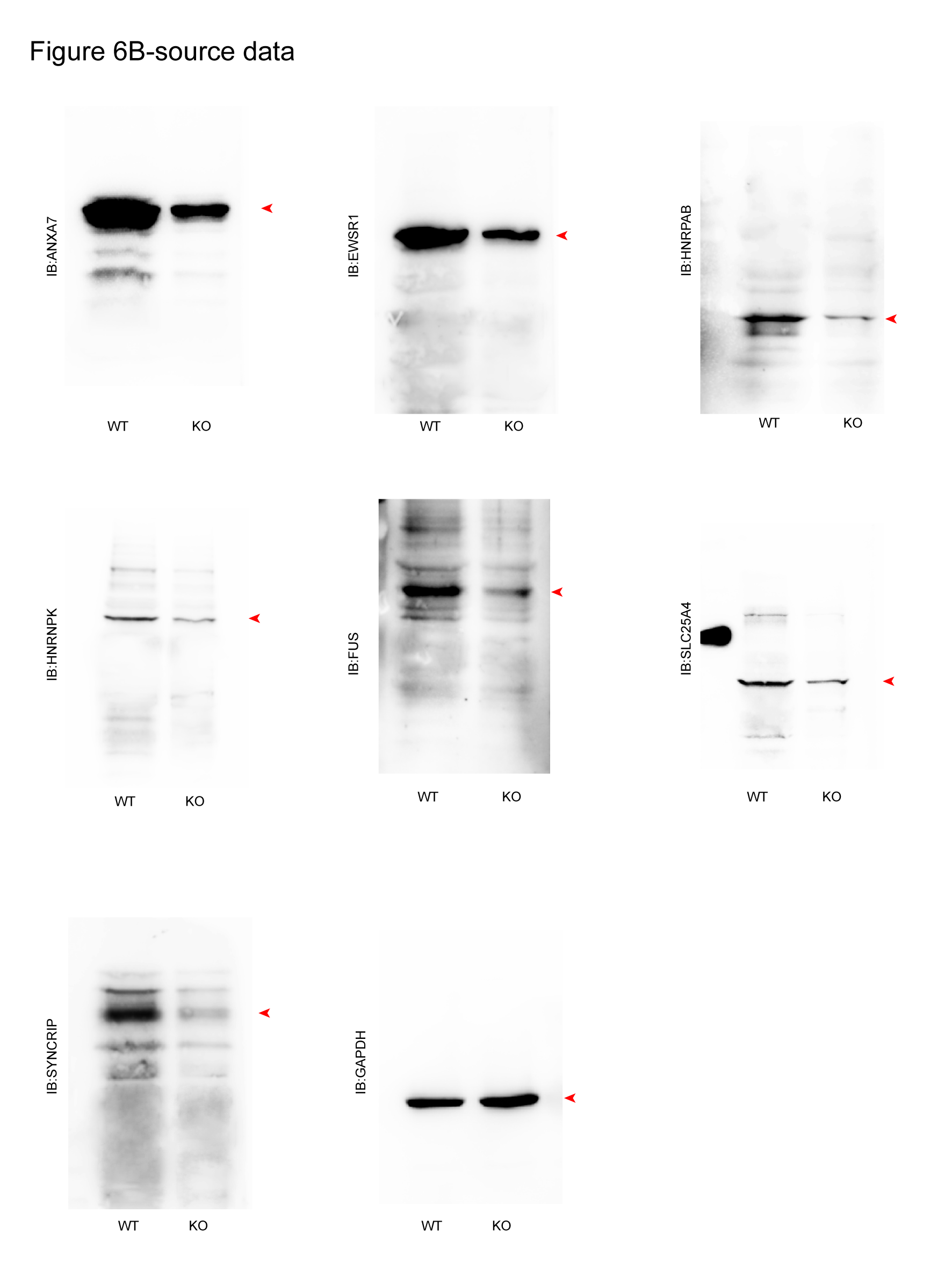

### Figure 6D-source data.tif

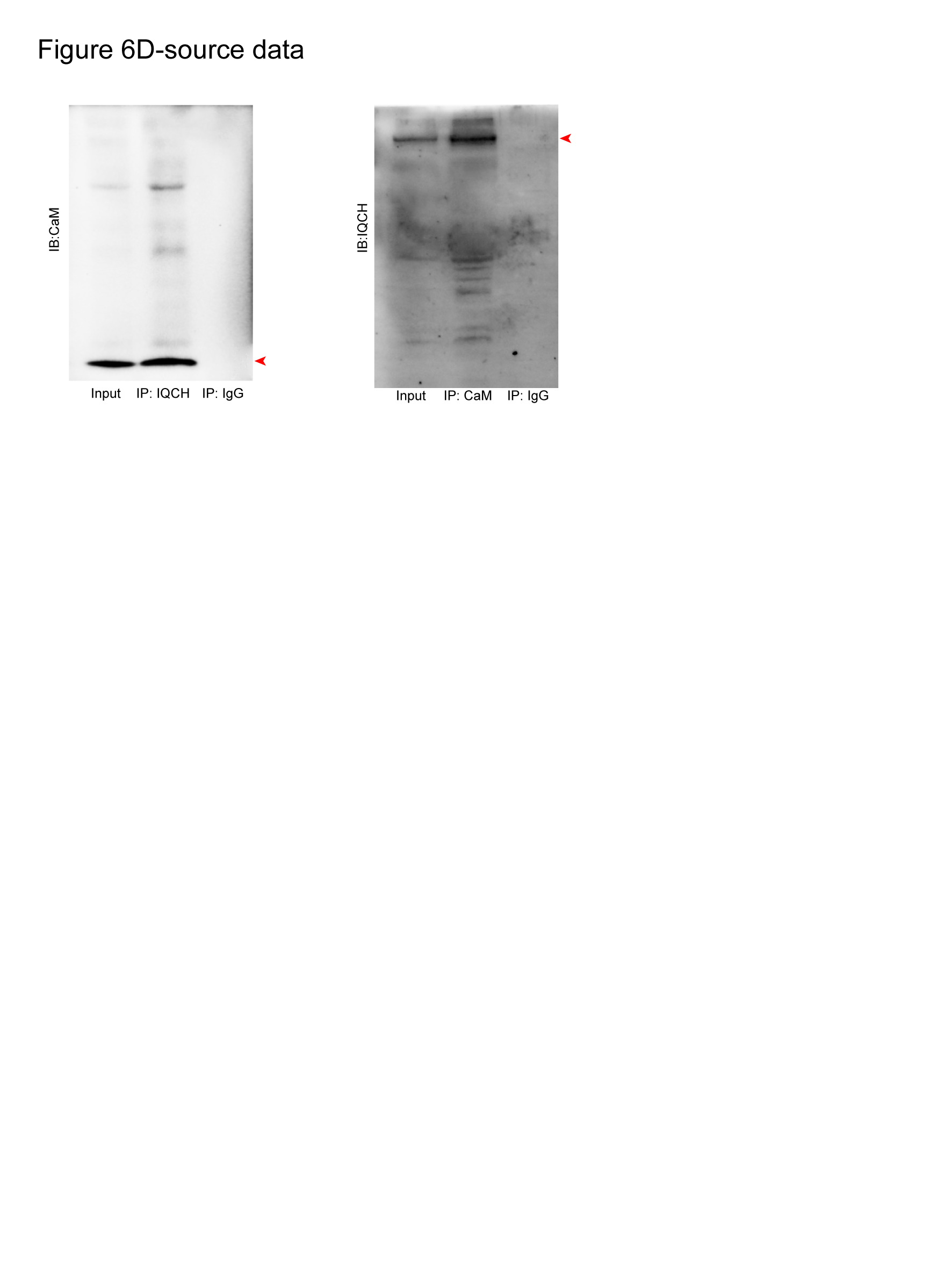

### Figure 6E-source data.tif

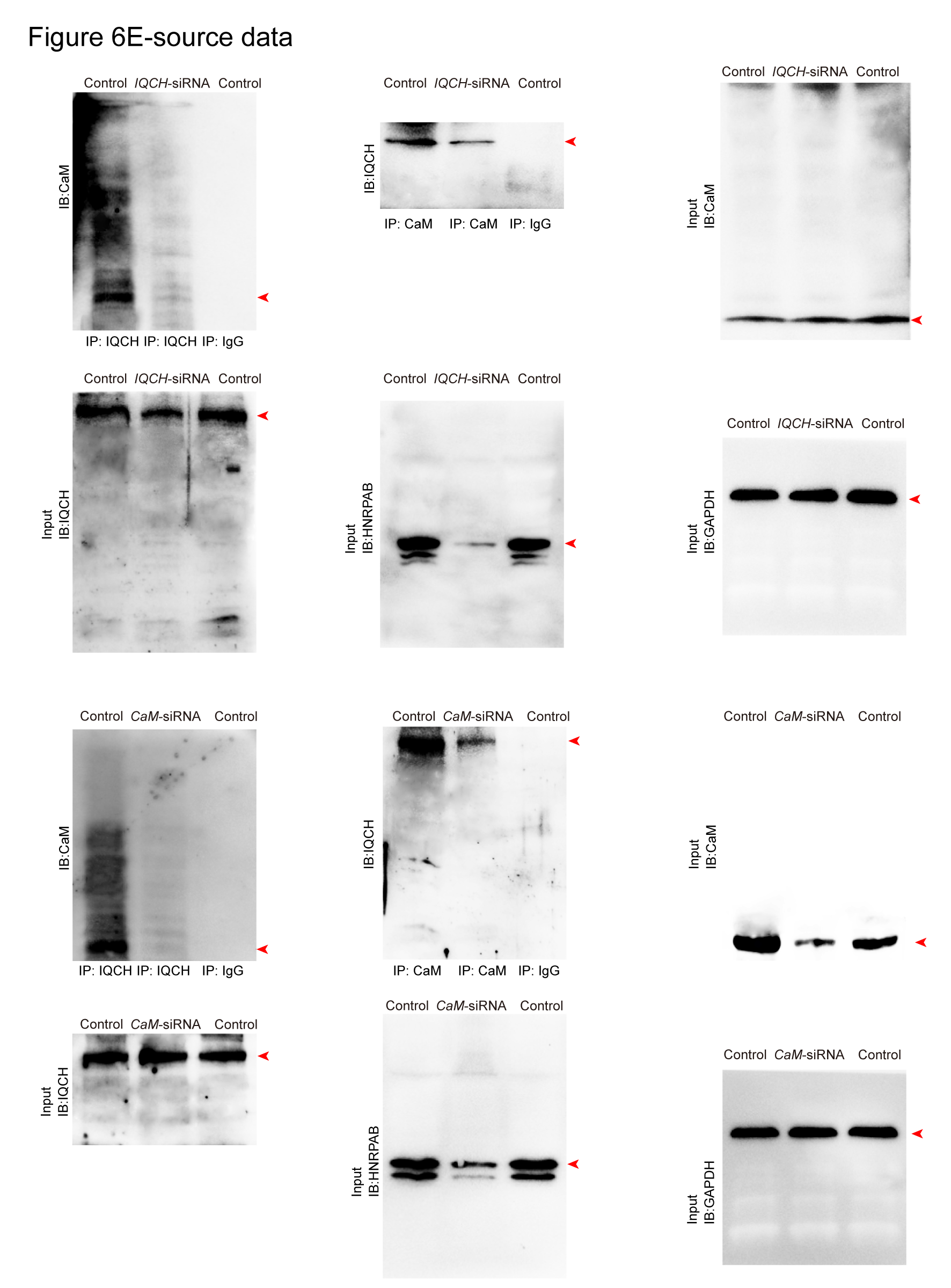

### Figure 6F-source data.tif

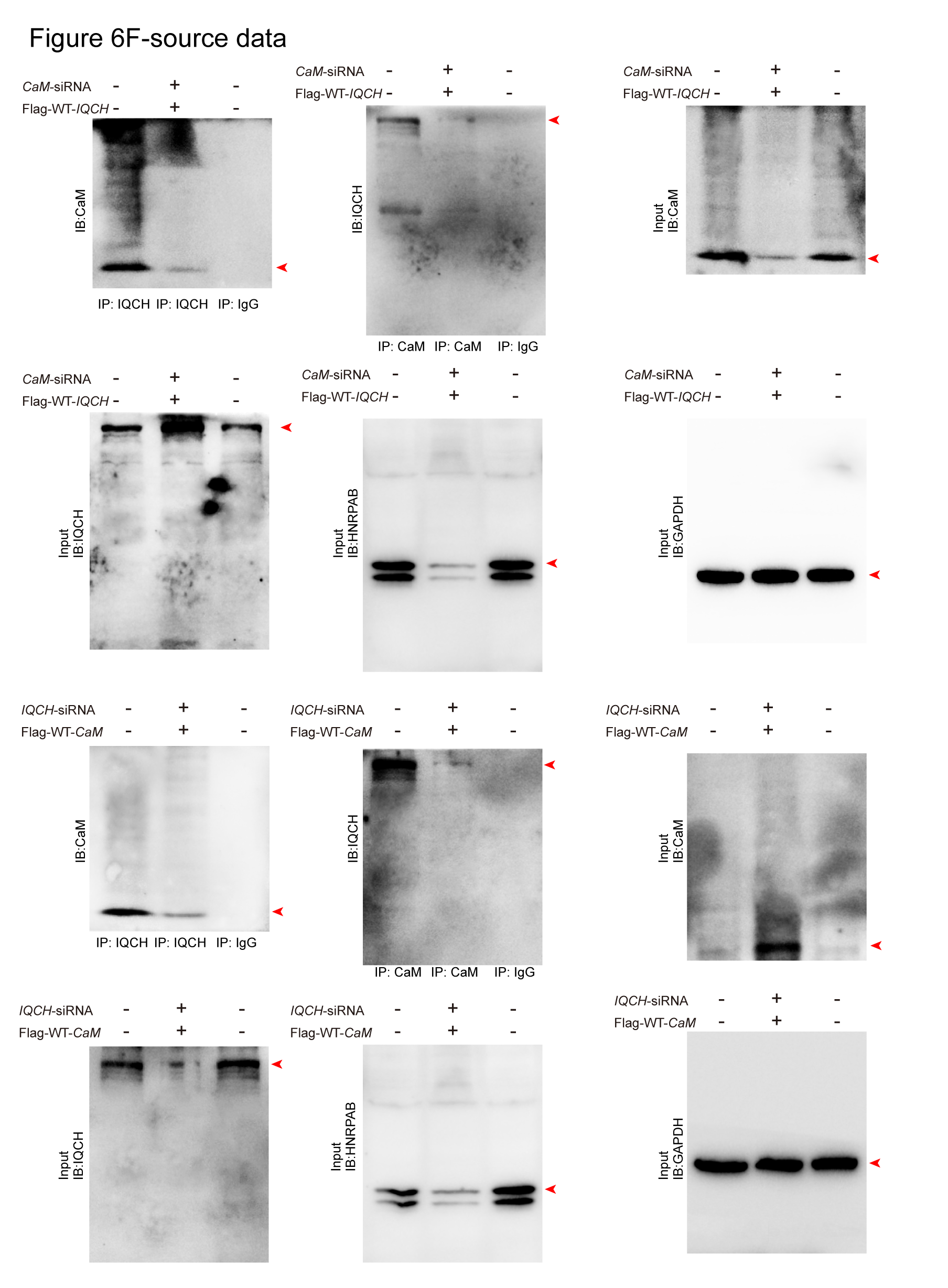

### Figure 6G-source data.tif

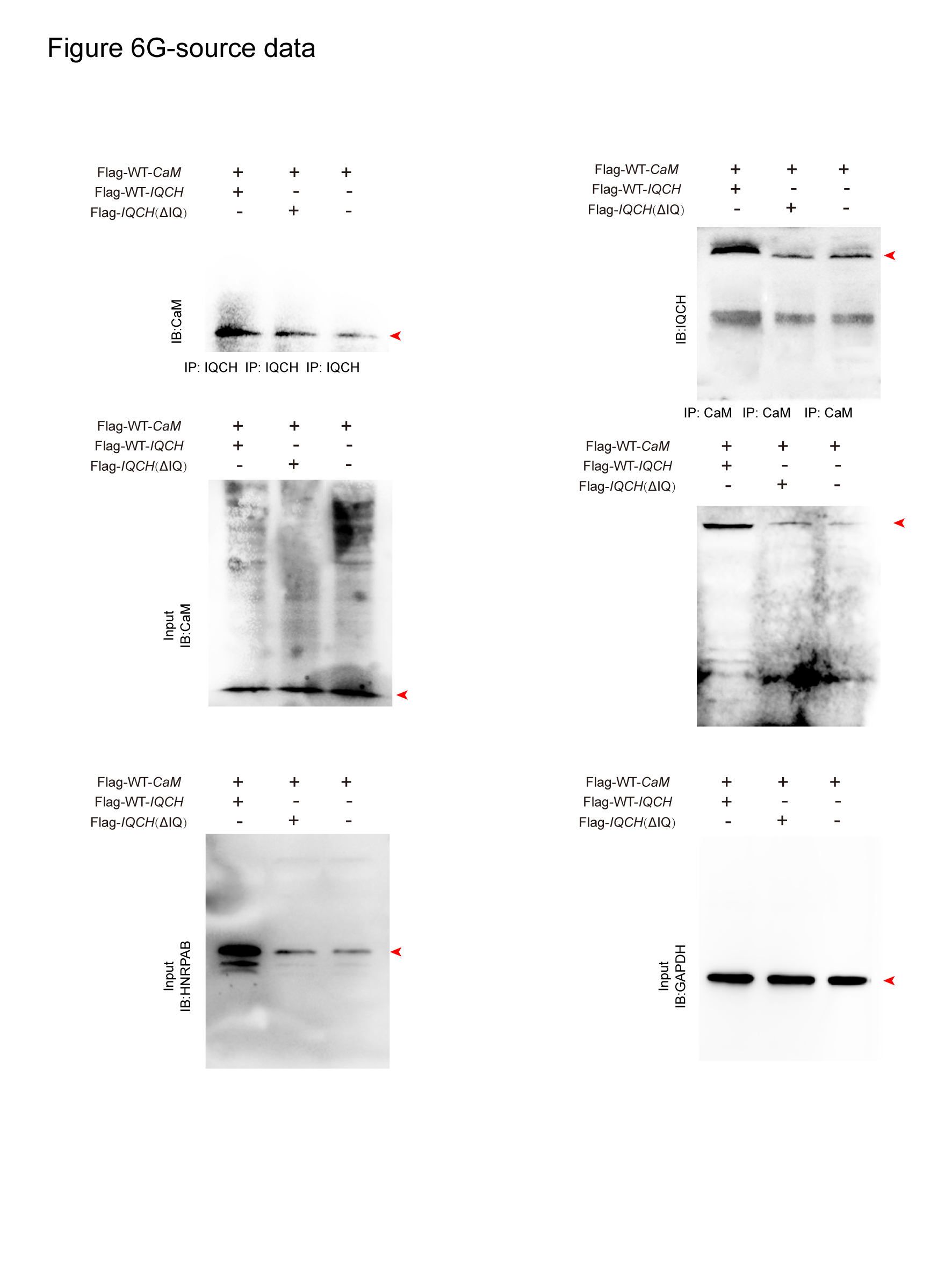

### Supplemental Data 1

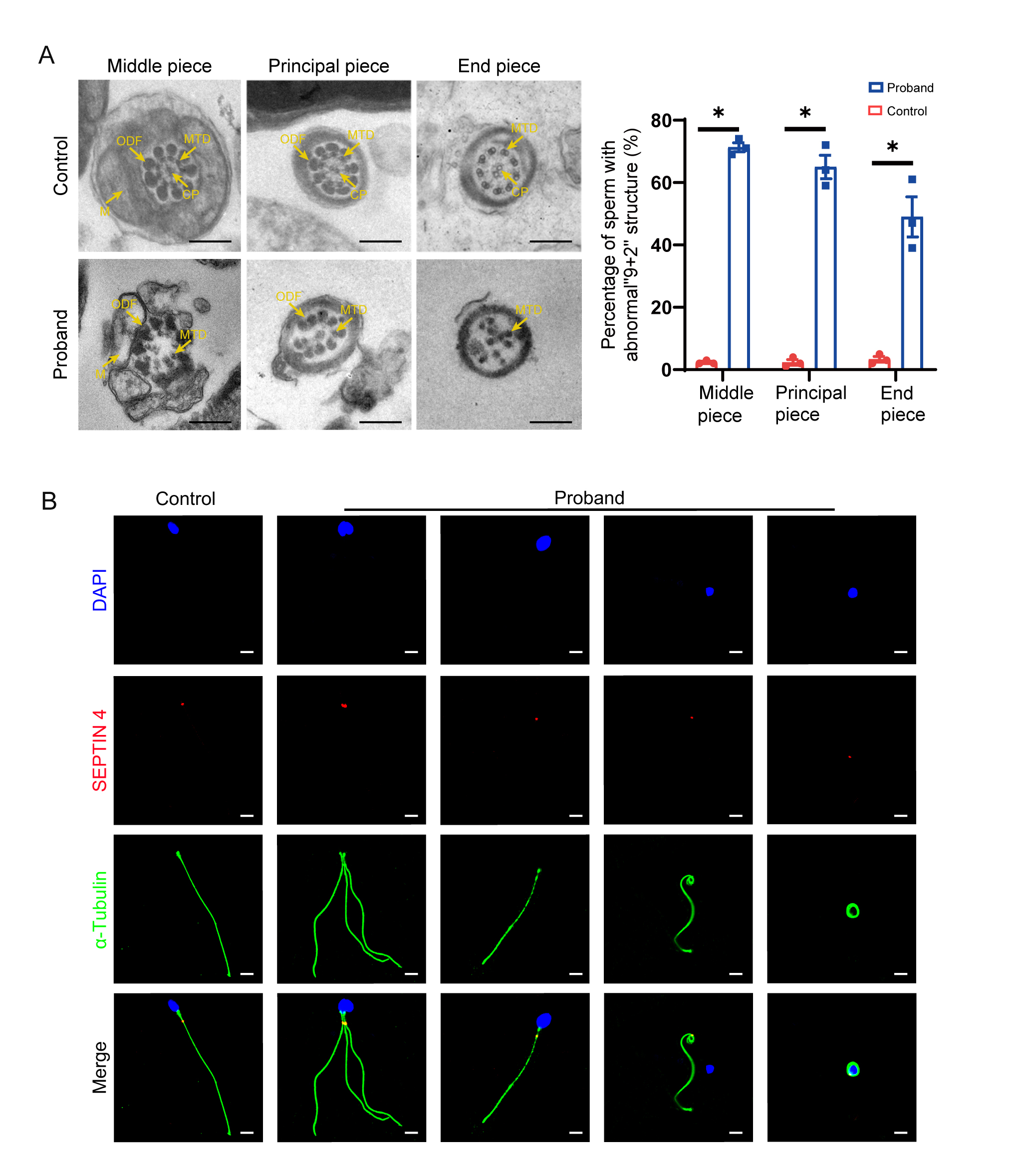

### Supplemental Data 2

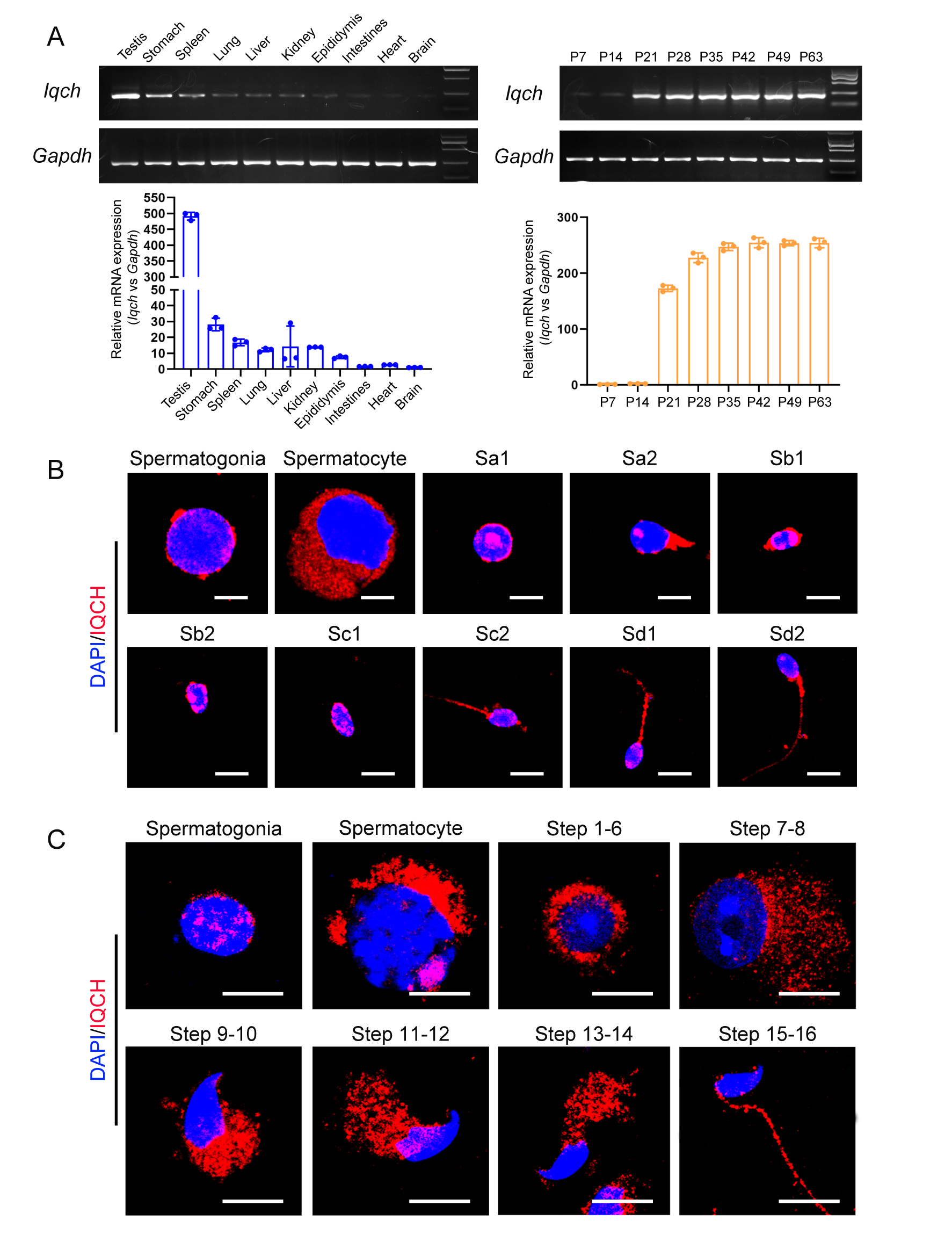

### Supplemental Data 3

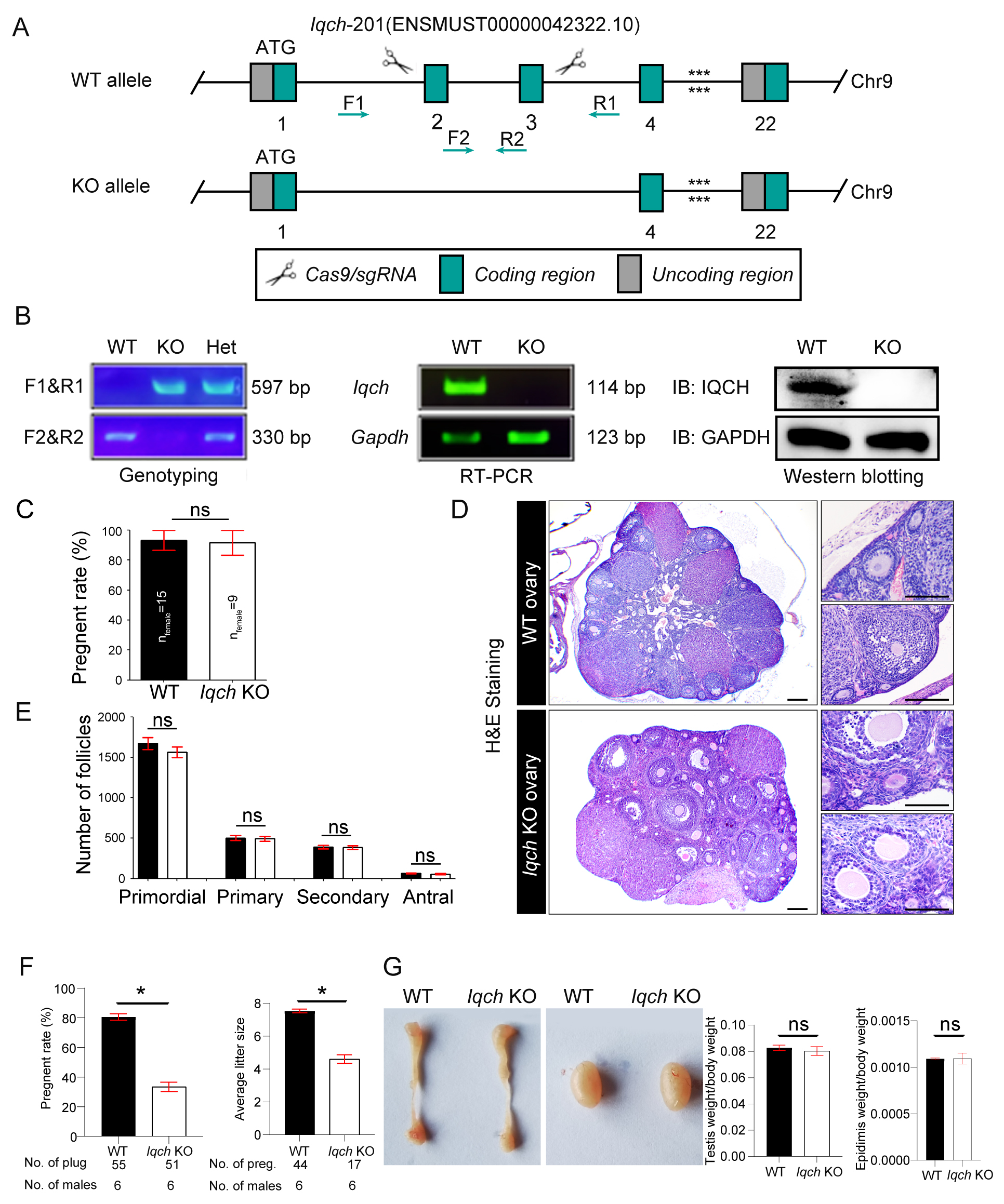

### Supplemental Data 4

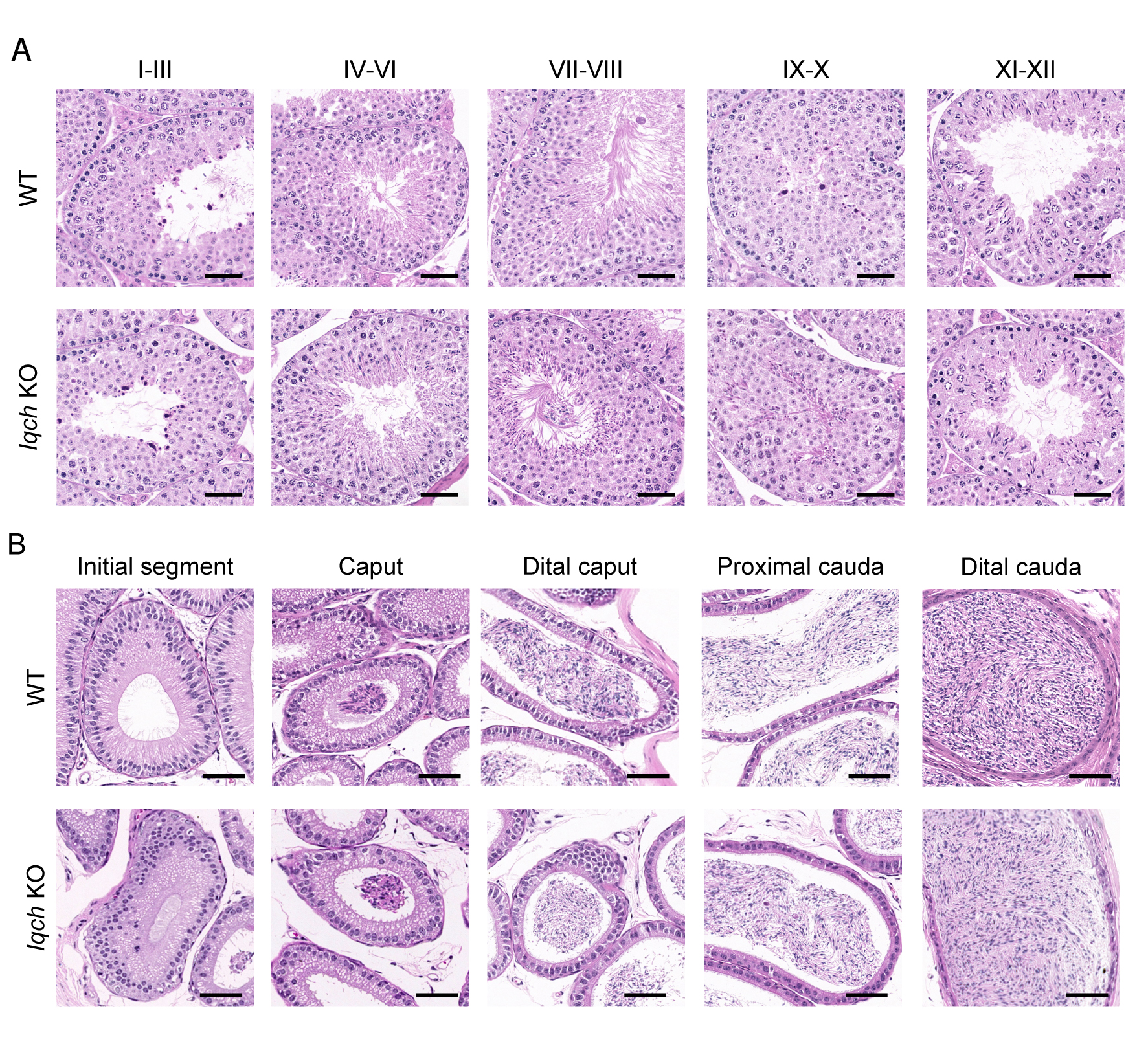

### Supplemental Data 5

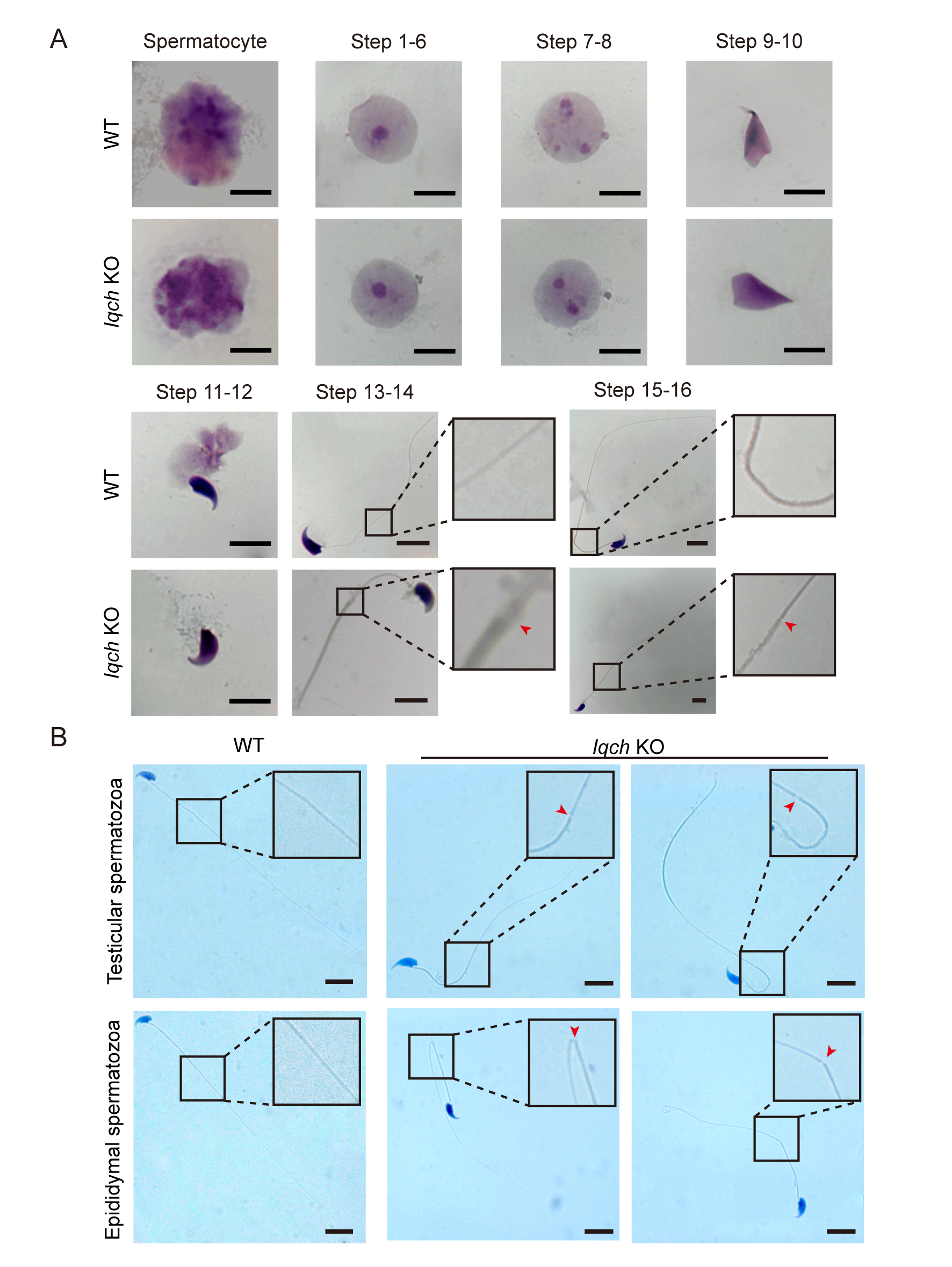

### Supplemental Data 6

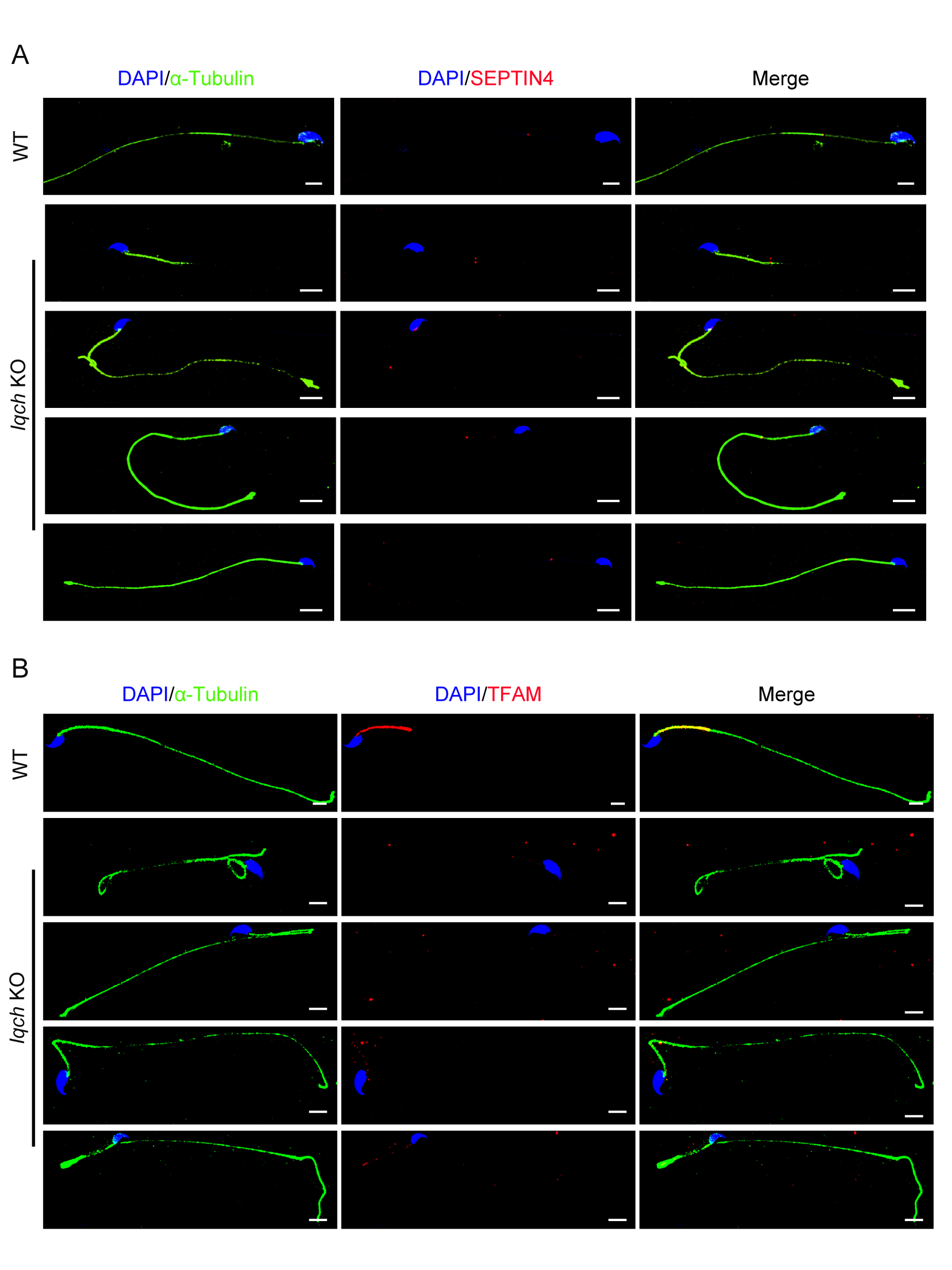

### Supplemental Data 7

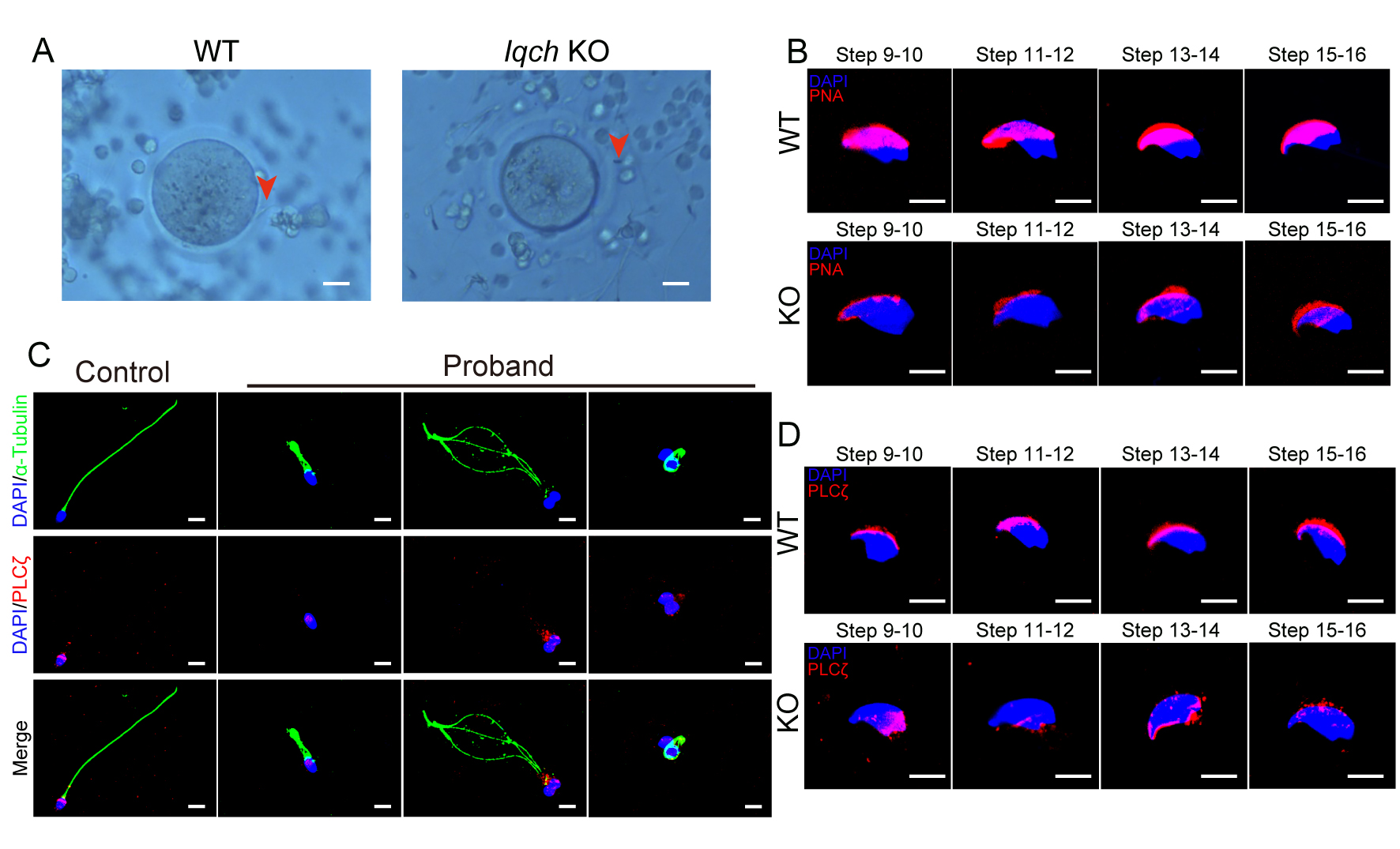

### Supplemental Data 8

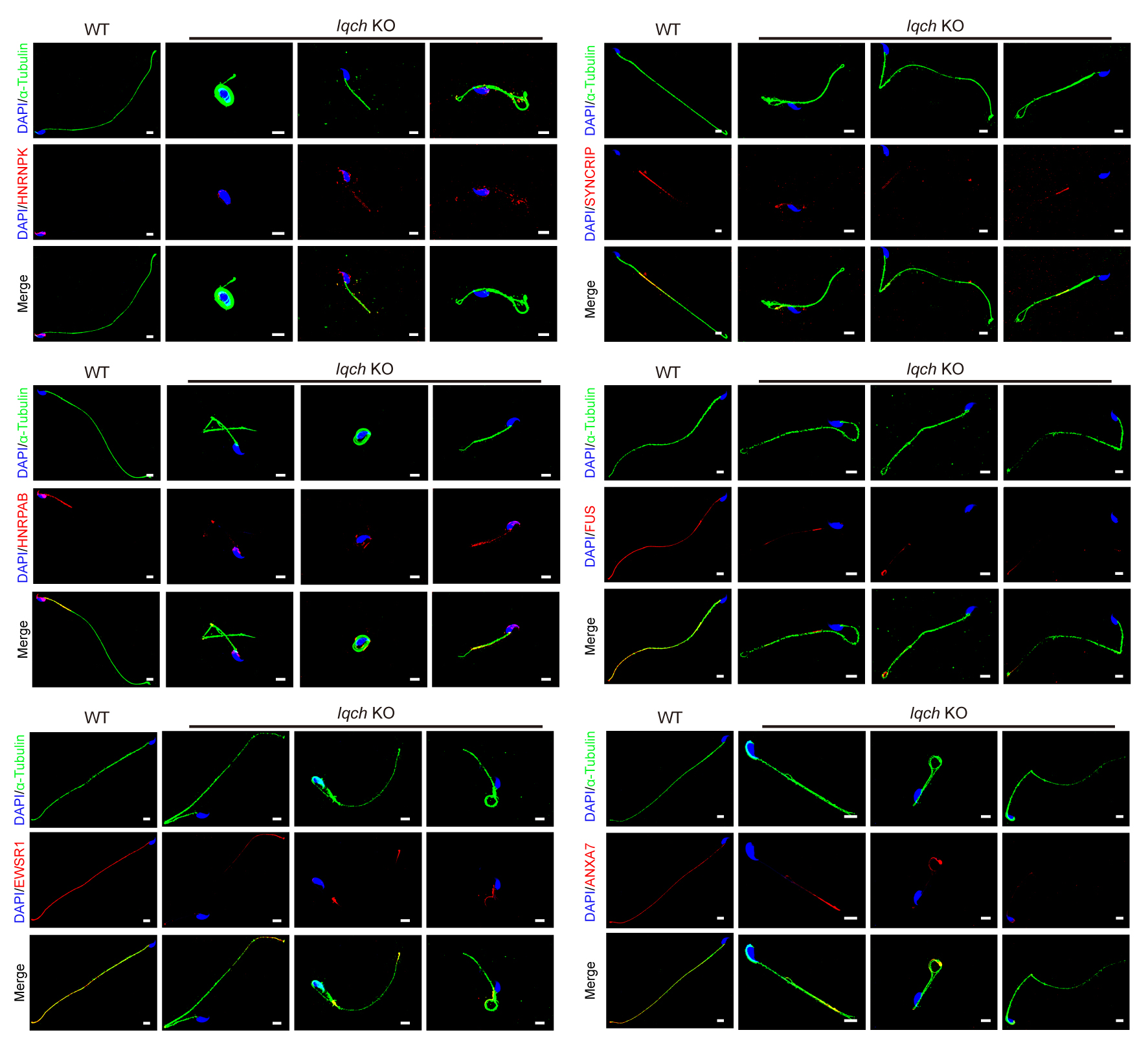

### Supplemental Data 9

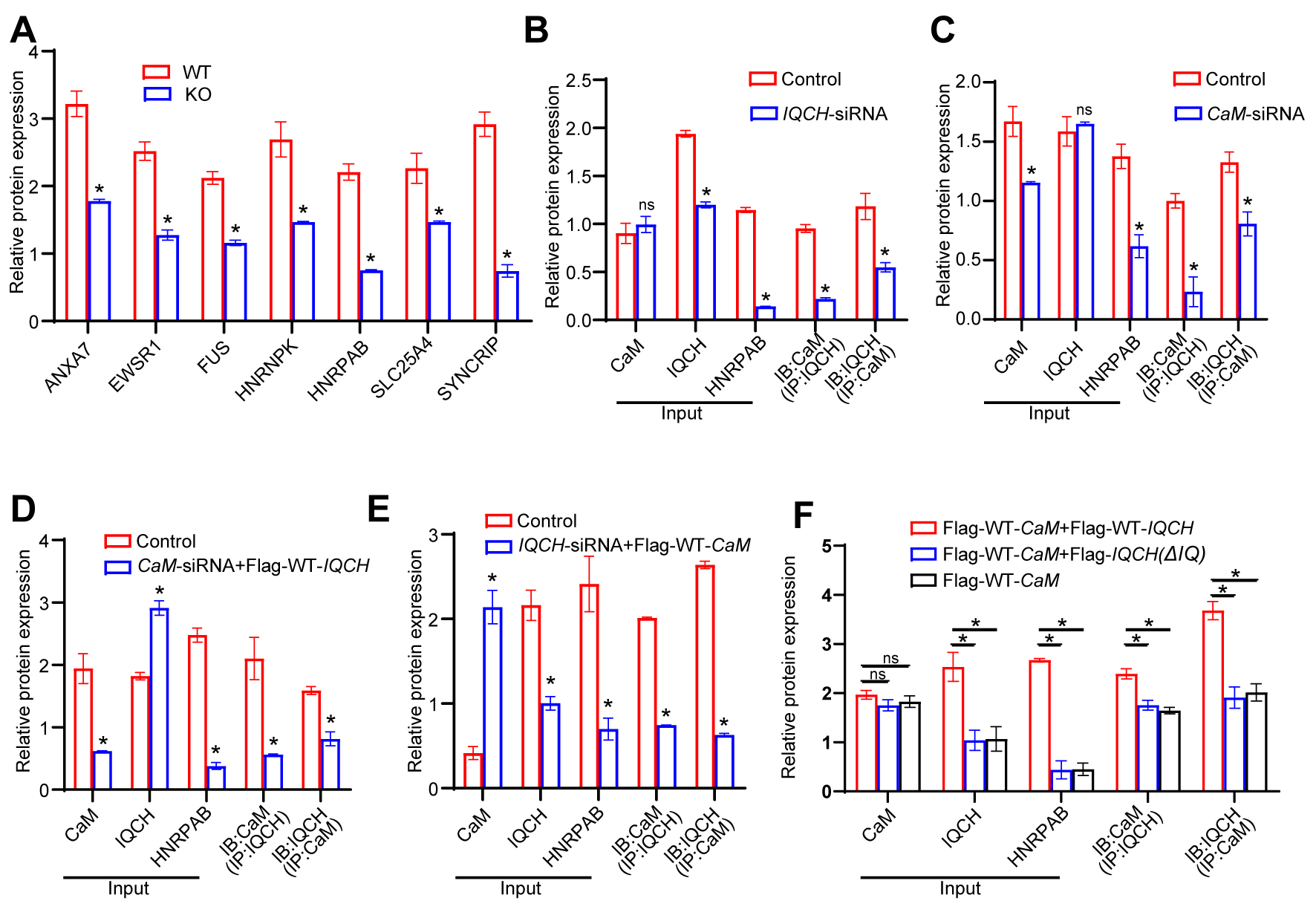

### Supplemental Data 10

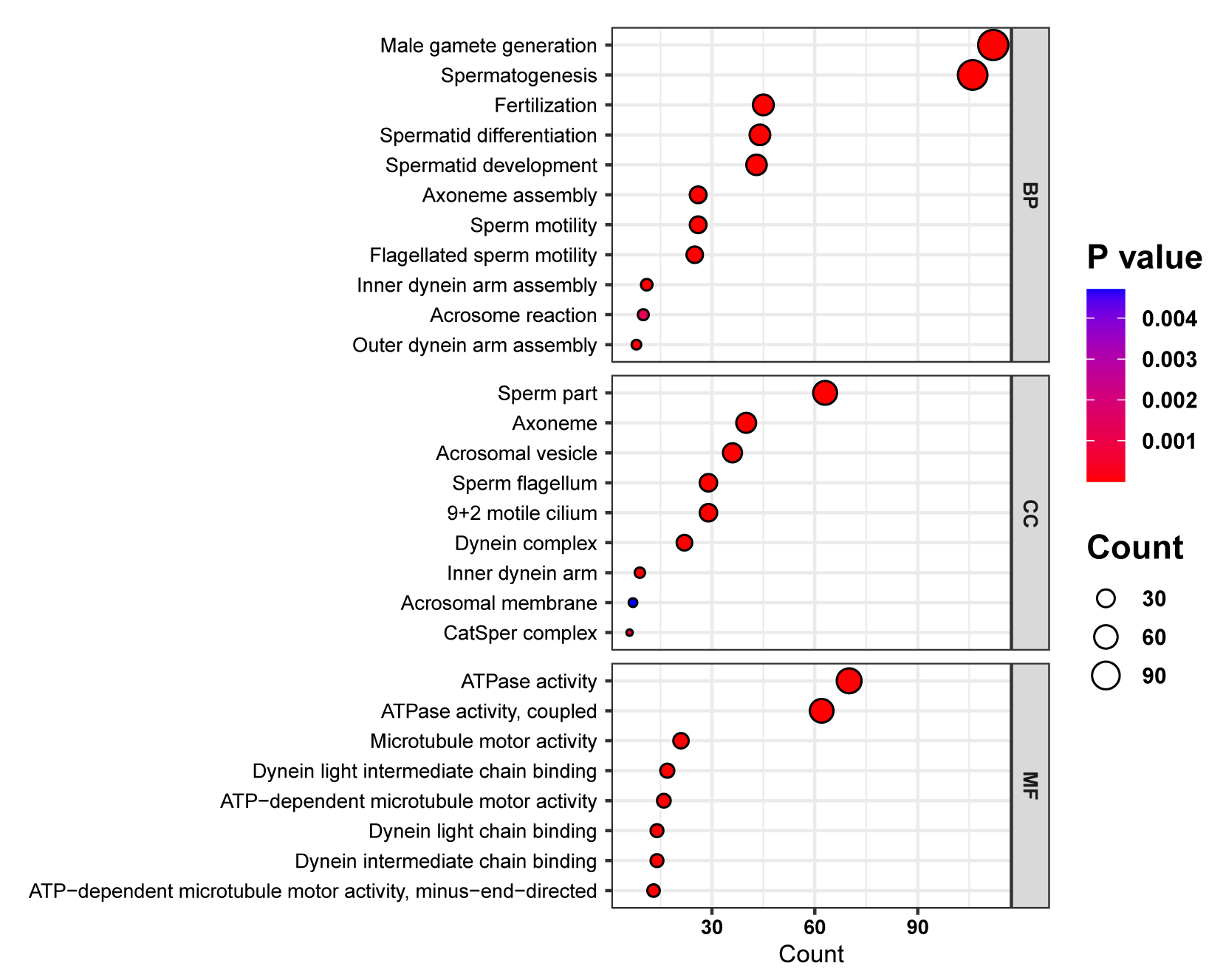

### Supplemental Data 11

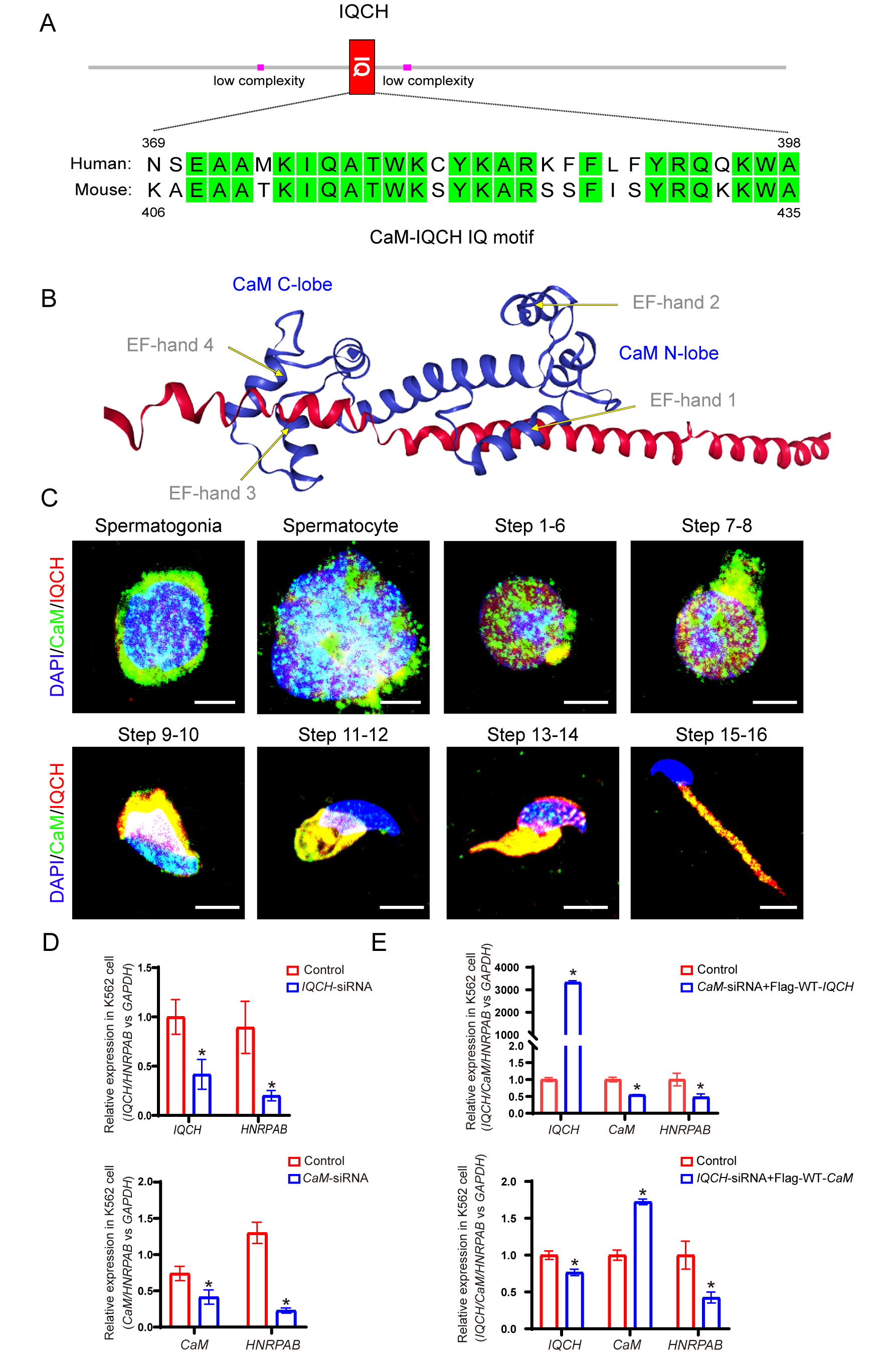
