## Supplementary material for "Deficiency of IQCH causes male infertility in humans and mice": Figure 1-source data 1

**Figure 1—source data 1.** Primers for Sanger sequencing and Minigene.

| **Variant** | **Sequence** |
| --- | --- |
| Sanger sequencing primer of *IQCH* | F 5' GACATGGCAGATGCAGAATCTG 3' |
|  | R 5' TGAATTTAGAGCAAGTTCATCAGAGA 3' |
| Amplification primer of *IQCH* for minigene assay | F 5' TACGGGATCACCAGTCCGCCTTCCGGGTTCATGCCACTCTC 3' |
|  | R 5' TCACCAGATATCTGGCAAGAGCCTATGGCCCACACCCTAGGC 3' |
| Amplification primer of plasmid for minigene assay | F 5' TCTGAGTCACCTGGACAACC 3' |
|  | R 5' ATCTCAGTGGTATTTGTGAGC 3' |
