## Supplementary material for "Deficiency of IQCH causes male infertility in humans and mice": Figure 3-source data 1

**Figure 3—source data 1.** Primers for RT-PCR and sgRNA sequences

| **Variant** | **Sequence** |
| --- | --- |
| RT-PCR for *Iqch* | F 5’ ATGCAGTGATGTCCACCCAG 3’ |
|  | R 5’ GCGAGCAAAGGTCATGAGGA 3’ |
| *Iqch* KO genotyping (F1/R1) | F 5’ CAGCAGCCCATGCAATAATC 3’ |
|  | R 5’ TAGGTGCTAATGGACACAGGCTAG 3’ |
| *Iqch* KO genotyping (F2/R2) | F 5’ TTCTAAGGAAAGCACCCACTGG 3’ |
|  | R 5’ TGAACAGAGACAGCAAGTCTGAGC 3’ |
| sgRNAs for *Iqch* KO mice | F 5’GGTACAACAAATGCGCCCAC 3’; PAM: GGG |
|  | F 5’AAGGGAATATAGATTACGAG 3’; PAM: TGG |
