## Supplementary material for "Deficiency of IQCH causes male infertility in humans and mice": Figure 6-source data 1

**Figure 6—source data 1.** Primers for qPCR and siRNA sequences

| **Variant** | **Sequence** |
| --- | --- |
| RT-PCR primer of *Catsper1* | F 5’ CATTCCTATCAGCAGGACAGGG 3’ |
|  | R 5’ GGGTGGGACAAAGGTTCACT 3’ |
| RT-PCR primer of *Catsper2* | F 5’ CTGGGGAGGAGTGAAGCTCTG 3’ |
|  | R 5’ ACAAAGCAACAGCCTGACATT 3’ |
| RT-PCR primer of *Catsper3* | F 5’ GGTCACTCTGGACAACCAAGA 3’ |
|  | R 5’ CCAGCTACGGCTACCTCTAA 3’ |
| RT-PCR primer of *Ccdc39* | F 5’ CCGTTCCTCCCTGAAACATCT 3’ |
|  | R 5’ AGGGCTTGCTTTTCCTCTGAA 3’ |
| RT-PCR primer of *Ccdc40* | F 5’ GCAGCAGGAGAAGATGATCCG 3’ |
|  | R 5’ TGGTAGTGGAACTCTGTCTTGG 3’ |
| RT-PCR primer of *Ccdc65* | F 5’ GCCCCATTACCTTAGCTCCA 3’ |
|  | R 5’ ATCTTCCTCGGACAGGGGTG 3’ |
| RT-PCR primer of *Dnhd1* | F 5’ TGCCACCAGACAAGGTGAAT 3’ |
|  | R 5’ CCAACAGGTCAACGCCTTTC 3’ |
| RT-PCR primer of *Dnah8* | F 5’ AAACATCAGCCCAGAGGTCG 3’ |
|  | R 5’ AGTGTCTTCGGGTTCGTCAC 3’ |
| RT-PCR primer of *Lrrc6* | F 5’ CACCATGGGCCGAATCACA 3’ |
|  | R 5’ AAGTCCCGGCACCATTTGTC 3’ |
| RT-PCR primer of *IQCH* | F 5' CTGAAAACCACGACCCTGTC 3' |
|  | R 5' GTTCGCATTTACAGCAGCTC 3' |
| RT-PCR primer of *CAM* | F 5' GGGAACATCTGGGTTATGCC 3' |
|  | R 5' GTCCATAGTCCACGCAGAGT 3' |
| RT-PCR primer of *HNRPAB* | F 5' GAGAACGGACATGAGGCCGT 3' |
|  | F 5' CAGCTCAGGCCACCAACGAACAT 3' |
| siRNAs sequences of *IQCH* | F 5' CAGUUAAAGGAGAAAUUAACA 3' |
|  | F 5' UGUUAAUUUCUCCUUUAACUG 3' |
| siRNAs sequences of *CaM* | F 5' GAUGAAGAAGUUGAUGAAAUG 3' |
|  | F 5' CAUUUCAUCAACUUCUUCAUC 3' |
