## Supplementary material for "Deficiency of IQCH causes male infertility in humans and mice": Figure 6-source data 2

| **Antibody or dye (application and dilution ratio)** | **Source** | **Identifier** |
| --- | --- | --- |
| Anti-CAM(IF,1:50; WB, 1:200) | Santa Cruz Biotechnology | sc-137079 |
| Anti-IQCH(IF,1:50; WB, 1:200) | HuaAn Biotechnology | Customizedly produced |
| Anti-SEPTIN4(IF, 1:50) | Abclonal | A10238 |
| Anti-HNRPAB(IF, 1:50; WB, 1:500) | Abclonal | A17479 |
| Anti-HNRNPK(IF, 1:20; WB, 1:500) | Proteintech | 11426-1-AP |
| Anti-SYNCRIP(IF, 1:50; WB, 1:500) | Proteintech | 14024-1-AP |
| Anti-TFAM(IF, 1:50) | Proteintech | 22586-1-AP |
| Anti-EWSR1(IF,1:50; WB, 1:500) | Proteintech | 55191-1-AP |
| Anti-FUS(IF, 1:50; WB, 1:1000) | Proteintech | 11570-1-AP |
| Anti-ANXA7(IF, 1:50; WB, 1:1000) | Proteintech | 10154-2-AP |
| Anti-SLC25A4(IF, 1:50; WB, 1:500) | Signalway | 32484 |
| Alexa Fluor 488 (IF, 1:500) | Thermo Fisher | A21206 |
| Alexa Fluor 594 (IF, 1:500) | Thermo Fisher | A11005 |
| Anti-HNRPAB for RIP | Santa Cruz Biotechnology | sc-376411 |
| Anti-acetylated alpha Tubulin(IF, 1:50) | Santa Cruz Biotechnology | sc-23950 |

**Figure 6-source data 2. Overview of the antibodies or dyes used in this study.**
